# UPR^mt^ scales mitochondrial network expansion with protein synthesis via mitochondrial import

**DOI:** 10.1101/2020.08.12.248161

**Authors:** Tomer Shpilka, YunGuang Du, Qiyun Yang, Andrew Melber, Nandhitha U. Naresh, Joshua Lavelle, Pengpeng Liu, Hilla Weidberg, Rui Li, Jun Yu, Lihua Julie Zhu, Lara Strittmatter, Cole M. Haynes

**Affiliations:** Department of Molecular, Cell and Cancer Biology, University of Massachusetts Medical School, Worcester, MA 01605, USA; Department of Cellular and Physiological Sciences, Life Sciences Institute, University of British Columbia, Vancouver, BC V6T 1Z3, Canada; Electron Microscopy Core, University of Massachusetts Medical School, Worcester, MA 01605, USA

## Abstract

As organisms develop, individual cells generate mitochondria to fulfill physiologic requirements. However, it remains unknown how mitochondrial network expansion is scaled to cell growth and impacted by environmental cues. The mitochondrial unfolded protein response (UPR^mt^) is a signaling pathway mediated by the transcription factor ATFS-1 which harbors a mitochondrial targeting sequence (MTS)^1^. Here, we demonstrate that ATFS-1 mediates an adaptable mitochondrial expansion program that is active throughout normal development. Developmental mitochondrial network expansion required the relatively inefficient MTS^2^ in ATFS-1, which allowed the transcription factor to be responsive to parameters that impact protein import capacity of the entire mitochondrial network. Increasing the strength of the ATFS-1 MTS impaired UPR^mt^ activity throughout development due to increased accumulation within mitochondria. The insulin-like signaling-TORC1^3^ and AMPK pathways affected UPR^mt^ activation^4,5^ in a manner that correlated with protein synthesis. Manipulation to increase protein synthesis caused UPR^mt^ activation. Alternatively, S6 kinase inhibition had the opposite effect due to increased mitochondrial accumulation of ATFS-1. However, ATFS-1 with a dysfunctional MTS^6^ constitutively increased UPR^mt^ activity independent of TORC1 function. Lastly, expression of a single protein with a strong MTS, was sufficient to expand the muscle cell mitochondrial network in an ATFS-1-dependent manner. We propose that mitochondrial network expansion during development is an emergent property of the synthesis of highly expressed mitochondrial proteins that exclude ATFS-1 from mitochondrial import, causing UPR^mt^ activation. Mitochondrial network expansion is attenuated once ATFS-1 can be imported.

## Main

The UPR^mt^ is a mitochondrial-to-nuclear signal transduction pathway regulated by the transcription factor ATFS-1 that is required for development and longevity during mitochondrial dysfunction^1,7,8^. Because ATFS-1 harbors a MTS and a nuclear localization sequence (NLS), its transcription activity is regulated by subcellular localization. If ATFS-1 is imported into mitochondria, it is degraded by the protease LONP-1^1^ (Fig. 1a). However, if a percentage of ATFS-1 fails to be imported into mitochondria, it traffics to the nucleus to activate a transcriptional response that includes mitochondrial chaperones^9,10^. Perturbations to OXPHOS or mitochondrial proteostasis activate the UPR^mt^ as both processes are required for mitochondrial protein import^11^.

**Fig 1.**
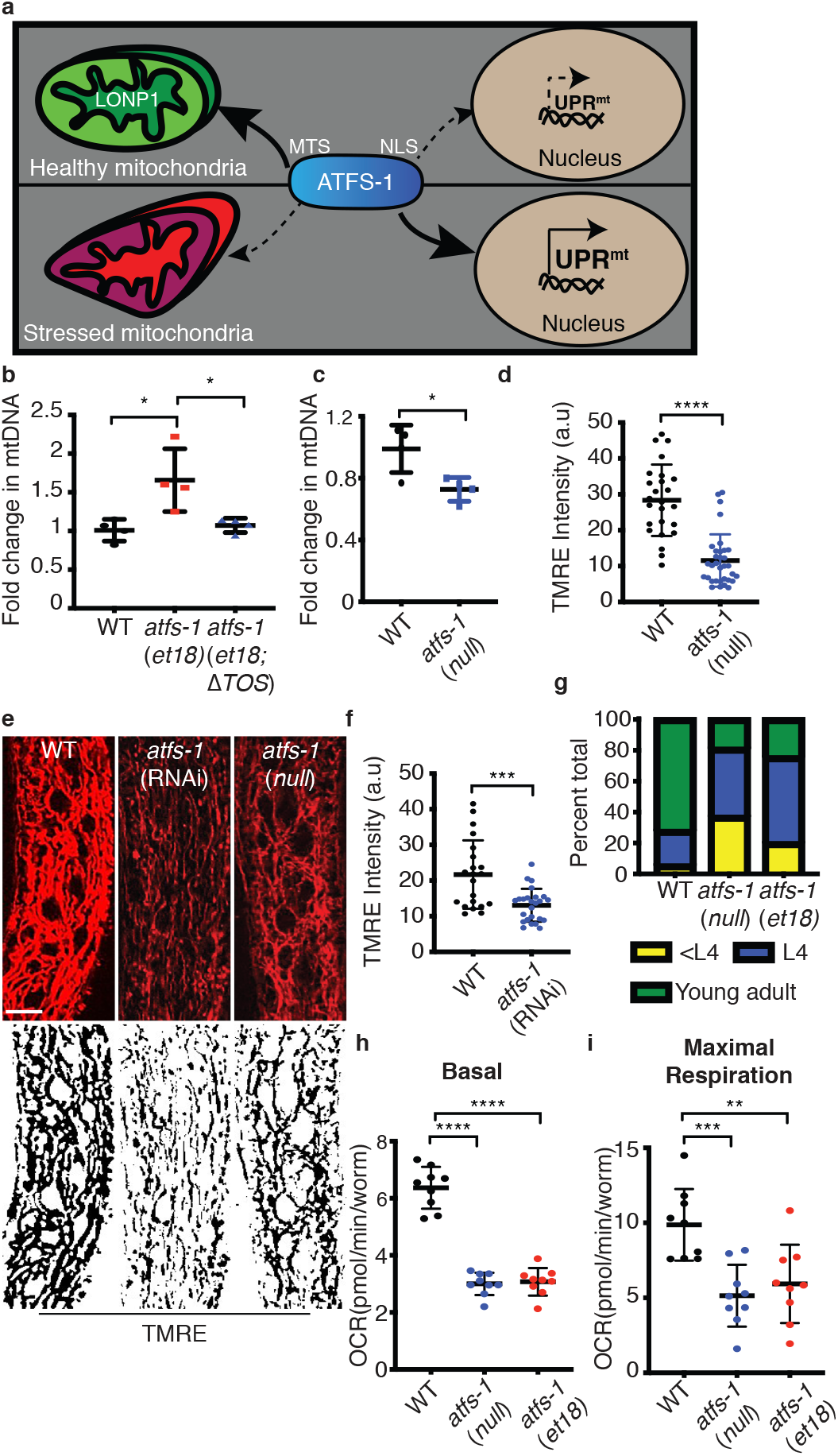
ATFS-1 regulates mitochondrial network expansion. **a.** Schematic of ATFS-1 regulation. **b.** Quantification of mtDNA in wildtype, *atfs-1(et18)* and *atfs-1(et18;△TOS)* as determined by qPCR. N=4. Error bars mean +/− s.d, *p<0.05 (Student’s *t*-test). **c.** Quantification of mtDNA in wildtype and *atfs-1(null)*. N=4. Error bars mean +/− s.d, *p<0.05 (Student’s *t*-test). **d.** Quantification of TMRE intensity in wildtype and *atfs-1(null)* worms. N=21(wildtype), N=33 *((atfs-1(null))*. Error bars mean +/− s.d, ***p<0.001, (Student’s *t*-test). **e.** TMRE staining of wildtype worms raised on control or *atfs-1*(RNAi) and *atfs-1(null)* worms. Skeleton-like binary backbone is presented (bottom). Scale bar 10μm. **f.** Quantification of TMRE intensity in wildtype and *atfs-1*(RNAi) worms. N=25. Error bars mean +/− s.d, ****p<0.0001 (Student’s *t*-test). **g.** Developmental stages of 3 day old wildtype, *atfs-1(et18)* or *atfs-1(null)* worms. N=546 (wildtype), N=597 atfs-1(*et18*) and 627 (*atfs-1(null)*). **h-i.** Oxygen consumption rates (OCR) in wildtype, *atfs-1(et18)* and *atfs-1(null)*. Basal respiration (**h**), maximal respiration (**i**). N=9. Error bars mean +/− s.d, **p<0.01,***p<0.001,****p<0.0001 (Student’s *t*-test).

### A role for ATFS-1 in mitochondrial network maintenance and expansion

We previously found that OXPHOS dysfunction due to deleterious mtDNA heteroplasmy caused an *atfs-1*-dependent expansion of the mitochondrial network that was observed only when mitophagy was impaired^12^. Similarly, OXPHOS dysfunction caused by mutations in the ubiquinone biogenesis gene *clk-1*, induced the UPR^mt^ and lead to an increase in mtDNA (Extended Data Fig. 1a) suggesting a role for the UPR^mt^ in mitochondrial biogenesis or network expansion.

*atfs-1(et18)* worms constitutively activate the UPR^mt^ due to an amino acid substitution in the MTS which impairs import into mitochondria even in the absence of mitochondrial stress^6^. Impressively, *atfs-1(et18)* worms harbored more mtDNAs relative to wildtype worms (Fig. 1b), suggesting that UPR^mt^ activation is sufficient to expand the mitochondrial network. Conversely, worms lacking the entire *atfs-1* open reading frame (*atfs-1(null)*)^13^ had reduced mtDNAs (Fig. 1c). Moreover, TMRE staining indicated that *atfs-1(null)* or *atfs-1* RNAi treated worms harbor fewer functional mitochondria relative to wildtype worms, in intestinal cells (Fig. 1d-f) suggesting the UPR^mt^ is actively involved in the maintenance and expansion of the mitochondrial network during development. Importantly, both *atfs-1(null)* and *atfs-1(et18)* caused developmental delays and impaired respiration (Fig. 1g-i). The reduction in respiratory capacity in the *atfs-1(et18)* strain was consistent with reduced TMRE staining in intestinal cells (Extended Data Fig. 1b-c). Thus, in the absence of the UPR^mt^, mtDNAs and functional mitochondrial are reduced, while continuous UPR^mt^ activation results in a partial expansion of the mitochondrial network that yields dysfunctional mitochondria.

### ATFS-1 mediates a mitochondrial expansion program during development

To elucidate the role of ATFS-1 in mitochondrial expansion and homeostasis, we compared transcriptional profiles of wildtype, *atfs-1(null)* and *atfs-1(et18*) worms during development in the absence of mitochondrial stress. Remarkably, *atfs-1(et18)* worms induced mitochondrial genes including the proteostasis components associated with the UPR^mt^ (Fig. 2a-d, Extended data Figure 2a, Supplementary table 1). Furthermore, over 50 genes required for mitochondrial ribosome function were upregulated, as were genes required for mtDNA replication, and cardiolipin biosynthesis pathway genes required for mitochondrial inner membrane synthesis. Lastly, genes required for both mitochondrial protein import and OXPHOS complex assembly were also upregulated.

**Fig 2.**
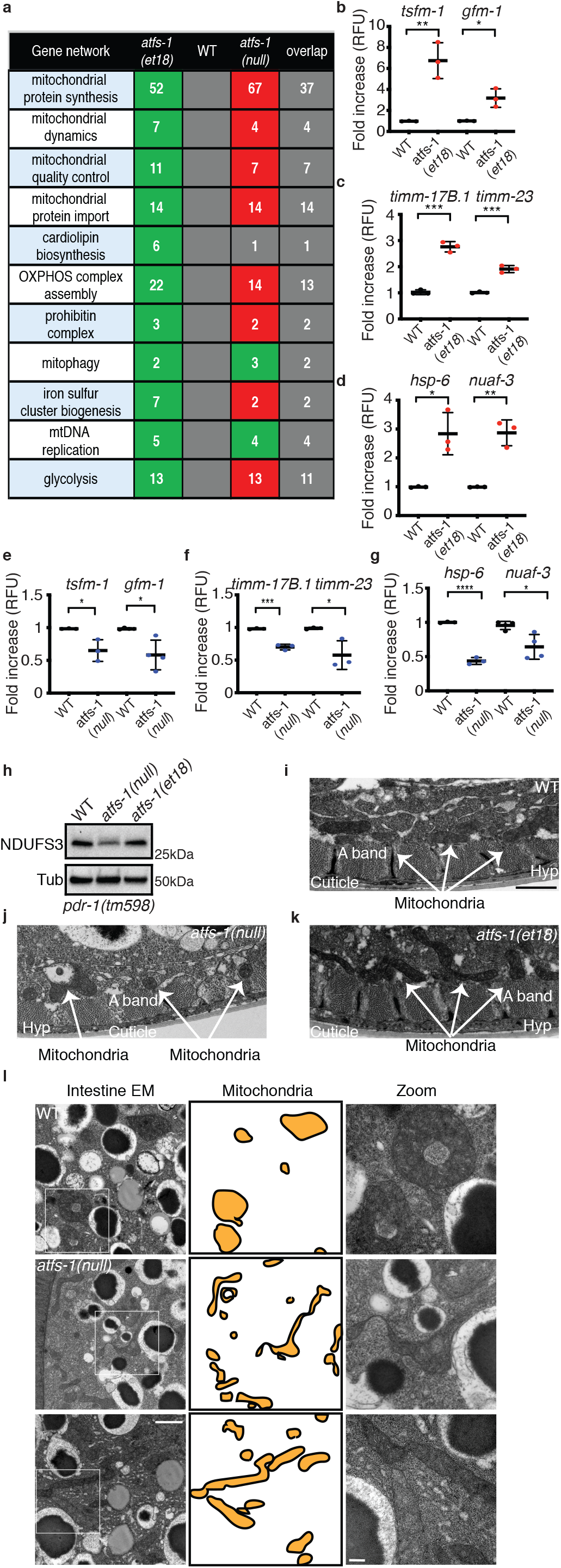
ATFS-1 mediates a mitochondrial network expansion program. **a**. Number of differentially regulated genes induced in *atfs-1*(*et18*) worms relative to wildtype (WT) or reduce in *atfs-1(null)* worms relative to wildtype. Green-upregulated, red-downregulated and the number of overlapping genes between *atfs-1(et18)* and *atfs-1(null)* worms are listed in the last column. **b-g**. Transcript levels of translation elongation factor mitochondrial 1 (*tsfm-1*) and G elongation factor mitochondrial 1 (*gfm-1*) (**b,e**), translocase of inner mitochondrial membrane 17B.1 (*timm-17B.1*) and translocase of inner mitochondrial membrane 23 (*timm-23*) (**c,f**), heat shock protein 6 (*hsp-6*) and NADH:ubiquinone oxidoreductase complex assembly factor 3 (*nuaf-3*) (**d,g**) as determined by qRT-PCR in wildtype and *atfs-1(et18)* (**b-d**) or in wildtype and *atfs-1(null)* (**e-g**). N=3 except for *gfm-1*(e) and *nuaf-3*(g) N=4. Error bars mean +/− s.d, *p<0.05, **p<0.01 ***p<0.001, ****p<0.001 (Student’s t-test). **h.** SDS-Page immunoblots of lysates from *pdr-1(tm598*), *atfs-1(null);pdr-1(tm598)* and *atfs-1(et18);pdr-1(tm598)* worms. NDUFS3 is a component of the NADH:Ubiquinone Oxidoreductase complex I and tubulin (Tub) was used as a loading control. **i-k.** Transmission electron microscopy of body wall muscle of wildtype (**i**), *atfs-1(null)* (**j**) and *atfs-1(et18)* worms (**k**). Scale bar 1 μm. **l.** Transmission electron microscopy of intestinal cells from wildtype, *atfs-1(null)* and *atfs-1(et18)* worms. Mitochondria are highlighted in the middle panel. Scale bar 1 μm (left) and 200nm (right).

Conversely, most of the mitochondrial genes induced in the *atfs-1(et18)* strain were downregulated in the *atfs-1(null)* worms compared to wildtype worms (Fig. 2a, 2e-g, Supplementary table 2-3), consistent with less mtDNA and TMRE staining in intestinal cells (Fig 1). Interestingly, the OXPHOS protein NDUFS3 was decreased in the *atfs-1(null),* while unaffected in *atfs-1(et18)* worms (Fig. 2h, Extended Data Fig. 2b). To exclude potential effects of mitochondrial degradation via mitophagy, these studies were performed in a strain lacking *pdr-1* (Parkin)^14^. Interestingly, multiple metabolic components including those of the TCA cycle and OXPHOS were repressed in *atfs-1(et18)* relative to wildtype worms (Supplementary table 4), consistent with UPR^mt^ limiting expression of highly expressed mitochondrial proteins^10^ and the reduced TMRE staining (Extended data Fig. 1c). Lastly, while many mitochondrial mRNAs were expressed at lower levels in *atfs-1(null)* worms, mtDNA replication and mitophagy components were upregulated suggesting an alternative stress response(s) is induced in the absence of *atfs-1* (Fig. 2a).

Because of the alterations in mtDNA levels, TMRE, and transcription of mitochondrial components, we visualized mitochondria via transmission electron microscopy. Impressively, mitochondria in *atfs-1(null)* worms were smaller and appeared defective in both intestine and muscle cells, along with pervasive muscle cell aberrations (Fig. 2i-l, Extended data Fig. 2c). In contrast, mitochondria in *atfs-1(et18)* were elongated, particularly visible in the intestine (Fig. 2i-l, Extended data Fig. 2c). Combined, our results suggest that ATFS-1 regulates a mitochondrial expansion program.

### A weak MTS regulates ATFS-1 and mitochondrial network expansion

We next sought to determine how ATFS-1 is regulated, or excluded from mitochondria, during development. Because ATFS-1 harbors a MTS along with a NLS, we have proposed that the UPR^mt^ is regulated by protein import capacity of the entire mitochondrial network^1^. ATFS-1 is predicted to have a relatively weak, or inefficient, MTS compared to other mitochondrial-targeted proteins such as mitochondrial chaperones and OXPHOS components^15,16^ (Fig. 3a). To compare the MTS strength of the OXPHOS protein ATP synthase subunit 9 (Su9) to ATFS-1, the amino-terminus of each was fused to GFP and expressed in HEK293T cells. As expected, both GFP-fusion proteins accumulated within mitochondria, but unlike Su9^(1-69)^::GFP, ATFS-1^(1-100)^::GFP fluorescence also accumulated within the cytosol, but to a lesser extent than that of ATFS-1^et18(1-100)^::GFP (Fig. 3b). Additionally, import of ATFS-1^(1-100)^::GFP was limited compared to Su9^(1-69)^::GFP in an *in vitro* import assay (Fig. 3d) consistent with ATFS-1 harboring a weak MTS.

**Fig 3.**
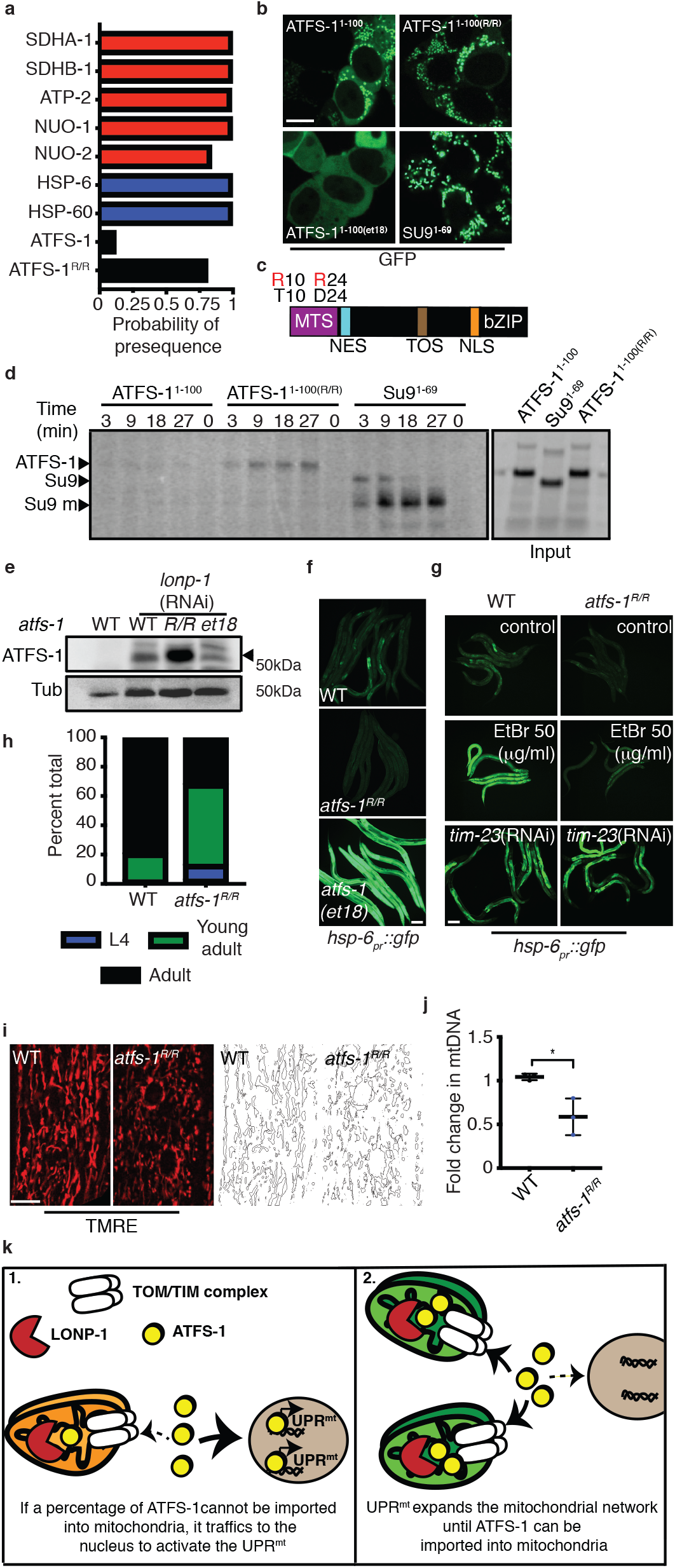
UPR^mt^ requires the weak MTS in ATFS-1. **a.** Mitochondrial targeting sequence probability prediction using MitoFates. OXPHOS proteins (red), mitochondrial chaperones (blue), ATFS-1 (black). **b.** HEK293T cells expressing ATFS-1^1-100^::GFP, ATFS-1^1-100(R/R)^::GFP, ATFS-1^1-100(et18)^::GFP or SU9^1-69^::GFP. Scale bar 10 μm. **c.** ATFS-1 schematic highlighting ATFS-1^R/R^ amino acid substitutions. **d.** In organelle import of radiolabeled ATFS-1^1-100^::GFP, ATFS-1^1-100(R/R)^::GFP and Su9^1-69^::GFP into isolated mitochondria. After the indicated time points, mitochondria were washed and analyzed by SDS-PAGE electrophoresis. ATFS-1, Su9, mature (m) are marked. **e.** SDS-Page immunoblots of lysates from wildtype, *atfs-1*^*R/R*^ and *atfs-1(et18)* worms raised on control or *lonp-1*(RNAi). ATFS-1 is marked (▸) and tubulin (Tub) was used as a loading control. **f.** *hsp-6pr::gfp* in wildtype, *atfs-1(et18)* and *atfs-1^R/R^* worms. Scale bar 0.1 mm. **g.** *hsp-6_pr_::gfp* in wildtype and *atfs-1^R/R^* worms raised on 50μg/ml EtBr, control or *timm-23*(RNAi). Scale bar 0.1 mm. **h.** Developmental stages of 3 day old wildtype or *atfs-1*^*R/R*^ worms. N=158 (wildtype) and N=256 (*atfs-1^R/R^*). **i.** TMRE staining of wildtype and *atfs-1*^*R/R*^ worms. Skeleton-like binary backbone is presented (right). Scale bar 10 μm. **j.** Quantification of mtDNA in wildtype and *atfs-1*^*R/R*^ worms as determined by qPCR. N=3. Error bars mean +/− s.d, *p<0.05 (Student’s *t*-test). **k.** Proposed model for ATFS-1 mediated mitochondria expansion.

We hypothesized that the inefficient MTS allows ATFS-1 import and UPR^mt^ activation to be sensitized to conditions that impact mitochondrial import capacity including mitochondrial stress, total mitochondria, and potentially the flux of other proteins into mitochondria. Thus, we sought to generate a worm strain expressing ATFS-1 with a stronger, or more efficient, MTS. Amino acid substitutions of T10 and D24 to arginine are predicted to increase MTS strength (Fig. 3a, 3c). Similar to Su9^(1-69)^::GFP, ATFS-1^R/R(1-100)^::GFP only accumulated within mitochondria and not in the cytosol in HEK293T cells (Fig. 3b). Furthermore, more ATFS-1^R/R(1-100)^::GFP accumulated within mitochondria than ATFS-1^(1-100)^::GFP in an *in vitro* import assay, consistent with increased MTS strength (Fig. 3d).

Via CRISPR-Cas9, mutations were introduced at the endogenous *atfs-*1 locus to generate ATFS-1^R/R^. We first examined accumulation of ATFS-1^R/R^ within mitochondria during normal development by raising worms on *lonp-1*(RNAi), which impairs ATFS-1 degradation within the matrix^1^. Strikingly, more ATFS-1^R/R^ accumulated within mitochondria compared to wildtype ATFS-1 or ATFS-1^et18^ during normal development (Fig. 3e). And, ATFS-1^R/R^ worms expressed less *hsp-6_pr_::gfp* relative to wildtype or *atfs-1(et18)* worms during normal development (Fig. 3f, Extended Data Fig. 3a) and had reduced expression of *hsp-6* and *timm-23* mRNAs (Extended Data Fig. 3b-c). ATFS-1^R/R^ also impaired UPR^mt^ activation caused by ethidium bromide (EtBr) exposure (Fig. 3g, Extended Data Fig. 3d). However, *timm-23*(RNAi) which impairs a component required for import of most proteins harboring amino-terminal MTSs^11^, caused UPR^mt^ activation in both ATFS-1^R/R^ and wildtype worms (Fig. 3g, Extended Data Fig. 3d) indicating ATFS-1^R/R^ is a functional transcription factor likely impaired due to increased mitochondrial accumulation. Similar to worms lacking *atfs-1*, worms expressing ATFS-1^R/R^ developed slower (Fig. 3h) and exhibited a perturbed and fragmented mitochondrial network in both intestine and muscle cells along with a reduction in mtDNA (Fig. 3i-j, Extended Data Fig. 3e-g).

Combined, these data suggest that ATFS-1 regulates a transcriptional program to expand mitochondrial biomass that is active throughout development and reliant on an inefficient MTS that confers sensitivity to conditions that impact mitochondrial import capacity. These findings suggest that during development a percentage of ATFS-1 cannot be imported into mitochondria of growing cells resulting in modest UPR^mt^ activation and mitochondrial network expansion (Fig. 3k).

### Interplay between protein synthesis, mitochondrial import, and ATFS-1

Previous screens for components required for UPR^mt^ activation identified multiple regulators of growth-related protein synthesis including the insulin-like receptor *daf-2*, *rheb-1*, *mTOR* (*let-363),* and *rsks-1* (S6 kinase)^4,5^. TORC1 regulates protein synthesis rates in response to diverse inputs including growth signals and cellular energetics^3^. Insulin-like signaling-TORC1 promotes protein synthesis by phosphorylating RSKS-1, which in turn, phosphorylates a ribosomal subunit^3^ (Extended Data Fig. 4a). Alternatively, the 5’ AMP-activated protein kinase (AMPK) limits TORC1 activity and protein synthesis when ATP levels are low^17,18^.

**Fig 4.**
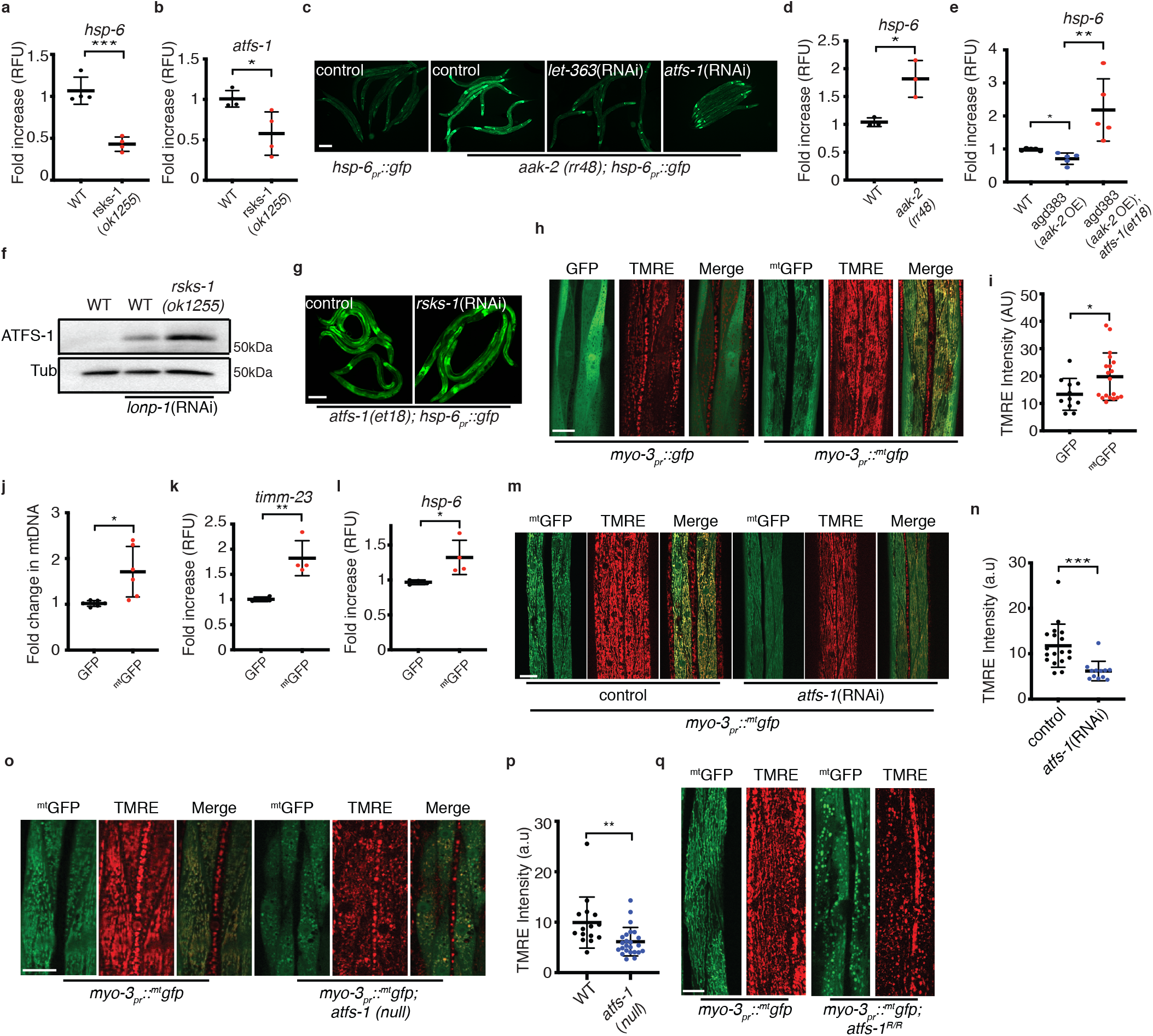
TORC1-mediated protein synthesis promotes ATFS-1 activation. **a-b**. Transcript levels of heat shock protein-6 (*hsp-6*) (**a**) and of activated transcription factor stress-1 (*atfs-1*) (**b**) as determined by qRT-PCR in wildtype and *rsks-1(ok1255)* worms. N=4. Error bars mean +/− s.d, *p<0.05, ***p<0.001 (Student’s t-test). **c.***hsp-6_pr_::gfp* and *aak-2(rr48);hsp-6_pr_::gfp* worms raised on control, *let-363* or *atfs-1* (RNAi). Scale bar 0.1 mm. **d-e.** Transcript levels of heat shock protein-6 (*hsp-6*) as determined by qRT-PCR in wildtype and *aak-2(rr48)* worms N=3 (**d**) and in wildtype, *agd383* and *agd383;atfs-1(et18)* strains N=5 (**e**). Error bars mean +/− s.d, *p<0.05, **p<0.01 (Student’s t-test). **f.** SDS-Page immunoblots of wildtype and *rsks-1(ok1255)* worms, raised on control or *lonp-1*(RNAi). Tubulin (Tub) was used as a loading control. **g.** *atfs-1(et18);hsp-6pr::gfp* worms raised on control or *rsks-1*(RNAi). Scale bar 0.1mm. **h.** TMRE staining (red) of worms expressing *myo-3pr::gfp* or *myo-3pr:: ^mt^gfp*. Scale bar 10μm. **i.** Quantification of TMRE intensity in muscle cells of worms expressing *myo-3pr::gfp* or *myo-3_pr_:: ^mt^gfp*. N=34 (*myo-3_pr_:: gfp*), N=24 (*myo-3_pr_:: ^mt^gfp)*. Error bars mean +/− s.d, *p<0.05 (Student’s *t*-test). **J.** Quantification of mtDNA in *myo-3_pr_:: gfp* and *myo-3_pr_:: ^mt^gfp* worms as determined by qPCR. N=6. Error bars mean +/− s.d, *p<0.05 (Student’s *t*-test). **k-l.** Transcript levels of translocase of inner mitochondrial membrane-23 (*timm-23*) (**k**) and of heat shock protein-6 (*hsp-6*) (**l**) as determined by qRT-PCR in *myo-3_pr_::gfp* and in *myo-3_pr_:: ^mt^gfp* worms. N=4. Error bars mean +/− s.d, *p<0.05, **p<0.01 (Student’s t-test). **m.** TMRE staining of worms expressing *myo-3pr:: ^mt^gfp* raised on control or *atfs-1*(RNAi). Scale bar 10 μm. **n.** Quantification of TMRE intensity in muscle cells of worms expressing *myo-3pr:: ^mt^gfp* raised on control or *atfs-1*(RNAi). N=18 (control), N=13 (*atfs-1*(RNAi)). Error bars mean +/− s.d, ***p<0.001 (Student’s *t*-test). **o.** TMRE staining of wildtype and *atfs-1*(*null*) worms expressing *myo-3_pr_:: ^mt^gfp*. Scale bar 10 μm. **o.** Quantification of TMRE intensity in muscle cells of wildtype, and *atfs-1(null)* worms. N=15 (wildtype), N=24 (*atfs-1(null)*). Error bars mean +/− s.d, **p<0.01 (Student’s *t*-test). **q.** TMRE staining of wildtype and *atfs-1^R/R^* worms expressing *myo-3_pr_:: ^mt^gfp*. Scale bar 10 μm.

As expected, DAF-2 inhibition impaired induction of *hsp-6_pr_::gfp* (Extended Data Fig. 4b)^19^, but *hsp-6* and *atfs-1* mRNAs were also reduced during normal development and during mitochondrial stress (Extended Data Fig. 4c-d). Inhibition of TORC1 components *rheb-1*, *raga-1, mTOR*, and *rsks-1* also reduced *hsp-6_pr_::gfp* (Extended Data Fig. 4e-g), as well as *hsp-6* and *atfs-1* mRNA levels, as did starvation (Fig. 4a-b, Extended Data Fig. 4h-j). Conversely, inhibition of AMPK, which increases TORC1 activity, resulted in increased *hsp-6* mRNA in an *atfs-1* and *mTOR*-dependent manner (Fig. 4c-d) while AMPK activation^20^ reduced *hsp-6* transcripts (Fig. 4e). Combined, our results indicate that TORC1 and RSKS-1 are required for UPR^mt^ during development and suggest that increased protein synthesis during development stimulates UPR^mt^ activity.

In mammals, TORC1 promotes protein synthesis by phosphorylating S6 kinase and 4EBP, which requires a TOR signaling (TOS) motif in each protein^21^. Interestingly, ATFS-1 also harbors a canonical TOS motif (-FEMDI-) (Fig. 3C), which we mutated at the endogenous locus to yield -AEMDI-(*atfs-1(ΔTOS)*). UPR^mt^ activation was attenuated in *atfs-1(ΔTOS)* worms relative to wildtype worms upon EtBr exposure (Extended Data Fig. 4k). Importantly, the TOS motif was also required for the increased *hsp-6_pr_::gfp* (Extended Data Fig. 4l) and mtDNA (Fig. 1b) in *atfs-1(et18)* worms suggesting the TOS motif promotes nuclear function of ATFS-1, similar to the TOS motif found in the transcription factor HIF-1α^22^. Interestingly, *hsp-6_pr_::gfp* induction caused by EtBr or complex III-deficiency (*isp-1(qm150)^23^*), was impaired further in *rsks-1(ok1255);atfs-1(ΔTOS)* worms relative to either *atfs-1(ΔTOS)* or *rsks-1(ok1255)* worms (Extended Data Fig. 4m-o), suggesting that RSKS-1 promotes UPR^mt^ activation independent of the TOS motif.

We next examined the effect of *rsks-1* inhibition on expression and trafficking of ATFS-1. One possibility is that *rsks-1* inhibition simply reduces synthesis of ATFS-1 limiting its nuclear transcription activity. Thus, we examined the mitochondrial accumulation of ATFS-1, by inhibiting LONP-1^1^. Interestingly, more ATFS-1 accumulated within mitochondria in *rsks-1(ok1255)* worms relative to wildtype worms (Fig. 4f). Thus, rather than reduced ATFS-1 expression, UPR^mt^ impairment in worms lacking RSKS-1 is due to mitochondrial accumulation of ATFS-1, similar to ATFS-1^R/R^ (Fig. 3e), which prevents trafficking to nuclei. Consistent with *rsks-1(ok1255)* impairing UPR^mt^ activation by increasing mitochondrial import capacity, *rsks-1*(RNAi) did not reduce *hsp-6_pr_::gfp* in *atfs-1(et18)* worms with an impaired MTS (Fig. 4g) or when treated with *timm-23*(RNAi) (Extended Data Fig. 5a). Moreover, activation of ATFS-1 persisted during starvation in *atfs-1(et18)* worms and *timm-23*(RNAi) treated worms (Extended Data Fig. 5b-c).

Because RSKS-1 is required for protein synthesis during cell growth, we hypothesized that the high rate import of proteins into mitochondria may cause UPR^mt^ activation and mitochondrial network expansion during normal development. As a test of this model, we sought to determine the impact of overexpressing a single protein with a relatively strong MTS on the mitochondrial network. The mitochondrial network was examined in muscle cells of worms expressing GFP or ^mt^GFP via the strong *myo-3* promoter^24^. Importantly, ^mt^GFP harbors a relatively strong MTS (aa 1-24) from the enzyme aspartate aminotransferase (AST) (Extended Data Fig. 5d), and the *myo-3* promoter is expressed throughout development^25^(Extended Data Fig. 5e). Relative to GFP, ^mt^GFP expression increased accumulation of functional mitochondria as determined by TMRE staining (Fig. 4h-i), mtDNA number (Fig 4j), and *hsp-6*, *timm-17* and *timm-23* mRNAs (Fig. 4k-l, Extended Data Fig. 5f) despite ^mt^GFP being expressed at a lower level than GFP (Extended Data Fig. 5g). Expansion of TMRE staining by ^mt^GFP was impaired by *atfs-1*(RNAi) (Fig. 4m-n), *atfs-1(null)* (Fig. 4o-p) and in ATFS-1^R/R^ worms (Fig. 4q), however ^mt^GFP transcription was not affected by the *atfs-1(null)* allele (Extended Data Fig. 5h).

To determine if the perturbed mitochondrial network was due to the inability of ATFS-1 to traffic to the nucleus and activate the UPR^mt^, we used CRISPR-Cas9 to generate impaired NLS in ATFS-1 (R426A). ATFS-1^(ΔNLS)^ accumulated within mitochondria similar to wildtype ATFS-1 (Extended Data Fig. 5i), but failed to induce *hsp-6_pr_::gfp* or endogenous *hsp-6* transcripts during mitochondrial stress elicited by knockdown of the mitochondrial protease SPG-7 (Extended Data Fig. 5j-k). Similar to ATFS-1^R/R^, or *atfs-1(null)* worms, ATFS-1^(ΔNLS)^ resulted in fragmented mitochondrial morphology in muscle cells indicating that the nuclear activity of ATFS-1 is essential for its function (Extended Data Fig. 5l). Combined, these finding indicate that expression of a single protein with a strong MTS is sufficient to expand the mitochondrial network in an *atfs-1*-dependent manner.

## Discussion

In summary, we have found that ATFS-1 regulates a mitochondrial expansion program that is active throughout normal development. Developmental mitochondrial expansion required the inefficient MTS of ATFS-1 and TORC1 activity suggesting an interplay between protein synthesis, mitochondrial protein import capacity, and nuclear activity of ATFS-1. Consistent with these findings, OXPHOS transcripts are among the most highly expressed mRNAs in worms (Extended Data Fig. 6a). And, *C. elegans* ribosome profiling data indicates that OXPHOS proteins are translated primarily during the early stages of worm development (L1,L2), and are reduced or absent by the L4 stage (Extended Data Fig. 6b). Interestingly, the ATFS-1 ribosome profile mirrors the OXPHOS profiles early in development and is also diminished at L4 (Extended Data Fig. 6c), consistent with the observation that the UPR^mt^ can only be activated by stress prior to the L4 stage^26^. Intriguingly, it was recently reported that mitochondrial metabolic proteins are prone to stalling within mitochondrial import channels under basal conditions in growing cells^27,28^, suggesting import or intra-mitochondrial protein processing can be overwhelmed during normal cell growth. We propose a model where the high levels of mitochondrial protein synthesis that occurs during development drives mitochondrial network expansion by excluding a percentage of ATFS-1 from mitochondrial import. And, network expansion continues until import is sufficient to import ATFS-1 and terminate the UPR^mt^. These findings are conceptually similar to the endoplasmic reticulum expansion that occurs in response to increased protein flux via the UPR^ER^, which is regulated by IRE1 and XBP1^29^.

We propose that as a function of the mitochondrial import flux or mitochondrial protein processing, ATFS-1 scales mitochondrial network expansion with cell growth.

## Methods

### Worms, plasmids and staining

The reporter strain *hsp-6_pr_::gfp* for visualizing UPR^mt^, the *myo-3_pr_::gfp* and the *myo-3_pr_::^mt^gfp* for visualization of mitochondrial mass and *atfs-1(null)* worms have been previously described^13,30,31^. The MTS in the *myo-3_pr_::^mt^gfp* is the first 24 amino acids from the enzyme aspartate aminotransferase from *Coturnix japonica* (1-MALLQSRLLLSAPRRAAATARASS-24) fused to GFP. The *atfs-1(et18)* strain was a gift from Marc Pilon. N2(wildtype), *isp-1(qm150)*, *rsks-1(ok1255)* and *daf-2(e1370)*, were obtained from the Caenorhabditis Genetics Center (Minneapolis, MN).

The *atfs-1^R/R^*(*cmh16*), the *atfs-1(ΔTOS)*(*cmh17*) and the *atfs-1(ΔNLS)(cmh18)* strains were generated via CRISPR-Cas9 in *hsp-6_pr_::gfp* worms as described^13^. The *atfs-1(ΔTOS)* was generated in both the wildtype worms as well as in the *atfs-1(et18)* strain. The crRNAs (IDT) were co-injected with purified Cas9 protein, tracrRNA (Dharmacon), repair templates (IDT) and the pRF4::rol-6(su1006) plasmid as described^32,33^. The crRNAs and repair templates used in this study are listed in Supplementary Table 5. The pRF4::rol-6 (su1006) plasmid was a gift from Craig Mello^34^. The ATFS-1^1-100^::GFP expressing plasmid was previously described^1^. The ATFS-1^1-100(R4C)^::GFP and the ATFS-1^1-100(T10R, D24R)^::GFP were generated by introducing mutations to yield the described amino acid substitutions in the ATFS-1^1-100^::GFP expressing plasmid. The subunit 9 of the F0-ATPase (SU9)^1-69^::GFP PQCXIP expression plasmid was a gift from Xuejun Jiang.

Worms were raised HT115 strain of *E. coli* and RNAi performed as described^35^. Ethidium bromide (EtBr) and TMRE experiments were performed by synchronizing and raising worms on plates previously soaked with M9 buffer containing EtBr or 2μM TMRE. Worms were analyzed at the L4 larvae stage except for EtBr treated worms that led to developmental arrest. EtBr treated worms were analyzed at the same time as the control.

### Protein analysis and antibodies

Synchronized worms were raised on plates with control(RNAi) or *lonp-1*(RNAi) to the L4 stage prior to harvesting. Whole worm lysate preparation was previously described^30^. Antibodies against α-tubulin were purchased from Calbiochem (CP06), GFP and for NDUFS3 from Abcam (ab6556 and ab14711 respectively). Antibodies for ATFS-1 were previously described^1^. Immunoblots were visualized using ChemiDoc XRS+ system (Bio-Rad). All western blot experiments were performed at least three times.

### mtDNA quantification

mtDNA quantification was performed using a qPCR-based method similar to previously described assays^36^. 20–30 worms were collected in 30 μl of lysis buffer (50 mM KCl, 10 mM Tris-HCl (pH 8.3), 2.5 mM MgCl_2_, 0.45% NP-40, 0.45% Tween 20, 0.01% gelatin, with freshly added 200 μg/ml proteinase K) and frozen at −80°C for 20 minutes prior to lysis at 65°C for 80 minutes. Relative quantification was used for determining the fold changes in mtDNA between samples. 1 μl of lysate was used in each triplicate qPCR reaction. qPCR was performed using the Thermo-Scientific SyBr Green Maxima Mix and the MyiQ2 Two-Color Real-Time PCR Detection System (Bio-Rad Laboratories). Primers that specifically amplify mtDNA are listed in Supplementary table 5. Primers that amplify a non-coding region near the nuclear-encoded *ges-1* gene were used as a control. mtDNA was harvested from synchronized worms at the L4 stage. All qPCR results have been repeated at least 3 times and performed in triplicates. A Student’s t-test was employed to determine the level of statistical significance.

### RNA isolation and qRT-PCR

RNA isolation and qRT-PCR analysis were previously described^12^. Worms were synchronized by bleaching, raised on HT115 *E. coli* and harvested at the L4 stage. Total RNA was extracted from frozen worm pellets using RNA STAT (Tel-Test) and 500 ng RNA was used for cDNA synthesis with qScript™ cDNA SuperMix (QuantaBio). qPCR was performed using iQ™ SYBR® Green Supermix (Bio-Rad Laboratories). qPCR primers are listed in Supplementary Table 5. All qPCR results were repeated at least 3 times and performed in triplicates. A Student’s t-test was employed to determine the level of statistical significance.

### Oxygen Consumption

Oxygen consumption was measured using a Seahorse XFe96 Analyzer at 25°C similar to that described previously^37^. In brief, L4 worms were transferred onto empty plates and allowed to completely digest the remaining bacteria for 1 hour, after which 10 worms were transferred into each well of a 96-well microplate containing 180 μl M9 buffer. Basal respiration was measured for a total of 30 minutes, in 6 minute intervals that included a 2 minute mix, a 2 minute time delay and a 2 minute measurement. To measure respiratory capacity, 15 μM FCCP was injected, the OCR (oxygen consumption rate) reading was allowed to stabilize for 6 minutes then measured for five consecutive intervals. Mitochondrial respiration was blocked by adding 40mM Sodium azide. Each measurement was considered one technical replicate.

### Cultured cells and imaging

HEK293T cells were transfected with 0.5 μg of the expression plasmids: SU9^1-69^::GFP with ATFS-1^1-100^::GFP, ATFS-1^1-100(R/R)^::GFP and ATFS-1^1-100(et18)^::GFP via Lipofectamine. The cells were imaged sixteen hours post transfection.

### RNA-sequencing and differential expression analysis

cDNA libraries were constructed with standard Illumina P5 and P7 adapter sequences. The cDNA libraries were run on an Illumina HiSEq2000 instrument with single-read 50-bp (SR50). RNA reads were then aligned to WBcel235/ce11 reference genome and differential gene expression analysis was performed with edgeR^38^. Differences in gene expression between *atfs-1(et18)* and *atfs-1(null)* compared to wildtype are listed in Supplementary Tables 6 and 7 respectively.

### Analysis of worm development

Worms were synchronized via bleaching and allowed to develop on HT115 bacteria plates for 3 days at 20^°^C. Developmental stage was quantified as a percentage of the total number of animals. Each experiment was preformed three times. For the comparison of wildtype and *atfs-1(null)* worms; N=162 (wildtype), and 282 (*atfs-1(null)*). For the comparison of wildtype to *atfs-1*^*R/R*^ worms; N=158 (wildtype) and N=256 (*atfs-1^R/R^*).

### Statistics

All experiments were performed at least three times yielding similar results and comprised of biological replicates. The sample size and statistical tests were chosen based on previous studies with similar methodologies and the data met the assumptions for each statistical test performed. No statistical method was used in deciding sample sizes. No blinded experiments were performed, and randomization was not used. For all figures, the mean ± standard deviation (s.d.) is represented unless otherwise noted.

### Microscopy

*C. elegans* were imaged using either a Zeiss AxioCam 506 mono camera mounted on a Zeiss Axio Imager Z2 microscope or a Zeiss AxioCam MRc camera mounted on a Zeiss SteREO Discovery.V12 stereoscope. Images with high magnification (63×) were obtained using the Zeiss ApoTome.2. Exposure times were the same in each experiment. Cell cultures were imaged with the Zeiss LSM800 microscope. All images are representatives of more than three images. Quantification of fluorescent intensity as well as creating binary skeleton-like structures were done with the Fiji software^39^.

### Gene set enrichment analysis

The OXPHOS gene set was downloaded from WormBase Ontology Browser^40^. mRNA abundance was measured and ranked by reads per kilobase per million reads (RPKM) from RNA-seq data. Pre-ranked gene set enrichment analysis was performed with GSEA3.0 software with ‘classical’ scoring^41^.

### Transmission Electron Microscopy

L4 larvae were transferred to 2.5% glutaraldehyde in 0.1 M Sodium Cacodylate buffer pH 7.2. for 10 min. The tail and head of each worm were dissected out and the main body was transferred to fresh 2.5% glutaraldehyde in 0.1 M Sodium Cacodylate buffer and kept at 4°C overnight. Samples were processed and analyzed at the University of Massachusetts Medical School Electron Microscopy core facility according to standard procedures. Briefly, the samples were rinsed three times in the same fixation buffer and post-fixed with 1% osmium tetroxide for 1h at room temperature. Samples were then washed three times with ddH_2_O for 10 minutes and then dehydrated through a graded ethanol series of 20% increments, before two changes in 100% ethanol. Samples were then infiltrated first with two changes of 100% Propylene Oxide and then with a 50%/50% propylene oxide/SPI-Pon 812 resin mixture. The following day, five changes of fresh 100% SPI-Pon 812 resin were performed before the samples were polymerized at 68°C in flat pre-filled embedding molds. The samples were then reoriented, and thin sections (approx. 70nm) were placed on copper support grids and contrasted with Lead citrate and Uranyl acetate. Sections were examined using a CM10 TEM with 100Kv accelerating voltage, and images were captured using a Gatan TEM CCD camera.

### Ribosome profiling data analysis

Ribosome profiling sequencing data was downloaded from the NCBI Sequence Read Archive (SRA) (http://www.ncbi.nlm.nih.gov/sra/) under accession number SRA055804. Data was analyzed as previously described^25^. Data analysis was done with the help of Unix-based software tools. First, the quality of raw sequencing reads was determined by FastQC^42^. Reads were then filtered according to quality via FASTQ for a mean PHRED quality score above 30^43^. Filtered reads were mapped to the *C. elegans* reference genome (Wormbase WS275) using BWA (version 0.7.5) and SAM files were converted into BAM files by SAMtools (version 0.1.19). Coverage data for specific genes (including 5’UTR, exons and 3’UTR) were calculated by SAMtools and coverage data for each gene was plotted using R^44^.

### Mitochondria isolation and *in vitro* protein import

Cells (budding yeast W303) were grown to logarithmic phase in YPD (1% yeast extract, 2% peptone, 2% glucose), collected by centrifugation and washed once with water. Cells were then resuspended in 0.1 M Tris pH 9.4, 10 mM DTT and incubated for 20 min at 30°C. Cell walls were disturbed by incubation in 1.2M sorbitol, 20mM K2HPO4 pH 7.4, 1% zymolyase for 1 h at 30°C. Dounce homogenization was used to lyse the cells in 0.6M sorbitol, 10mM Tris pH 7.4, 1mM EDTA, fatty acid free 0.2% BSA and 1mM PMSF. Mitochondria were then isolated by differential centrifugation as described previously^45^ and resuspended in SEM buffer (0.25M sucrose, 10mM MOPS KOH pH 7.2 and 1mM EDTA).

The coupled Transcription/Translation system (T7 Quick for PCR DNA, Promega) was used to express ATFS-1 from a PCR template. Precursor proteins (ATFS-1^1-100^::GFP, ATFS-1^1-100(R/R)^::GFP and Su9^1-69^::GFP) were synthesized in reticulocyte lysate in the presence of [35S]methionine (T7 Quick for PCR DNA, Promega). Import into isolated mitochondria was performed in import buffer (3 % (w/v) BSA, 250 mM sucrose, 80 mM KCl, 5 mM methionine, 5 mM MgCl2, 2 mM KH2PO4, 10 mM MOPS-KOH, pH 7.2, 4 mM NADH, 2 mM ATP, 5 mM creatine phosphate, 0.1 mg/ml creatine kinase) at 25°C. The import reaction was stopped on ice or by addition of AVO (8 μM antimycin A, 20 μM oligomycin, 1 μM valinomycin). To dissipate Δψ, AVO was added before the import experiment. Samples were treated with 25 μg/ml proteinase K for 15 min on ice, following by treatment with 2 mM PMSF for 5 min on ice. Mitochondrial were washed twice with SEM buffer and analyzed by electrophoresis on SDS-PAGE.

## Data availability

The data reported in this paper have been deposited in the Gene Expression Omnibus (GEO) database, https://www.ncbi.nlm.nih.gov/geo (accession no. GSE114951). Data also available from the corresponding author upon reasonable request.

## Acknowledgements

We thank W. Mair, M. Pilon and the *Caenorhabditis* Genetics Center for providing *C. elegans* strains (funded by NIH Office of Research 362 Infrastructure Programs (P40 OD010440), and the UMass Medical School Core facilities for deep sequencing and electron microscopy. This work was supported by HHMI, the Mallinckrodt Foundation, and National Institutes of Health grants (R01AG040061 and R01AG047182) to C.M.H. and (SI0OD021580) to L.S. The authors are solely responsible for the content.

## Author Contributions

T.S. and C.M.H. planned the experiments. T.S., A.M., N.U.N, Y.D., Q.Y. and J.L. generated worm strains. T.S., A.M., Y.D. and Q.Y. performed the western blots. T.S., J.L. and Y.D. performed the mitochondrial imaging, RNA-seq and UPR^mt^ regulation experiments. T.S. and Y.D. preformed seahorse experiments. J.Y., R.L. and L.J.Z. analyzed the sequencing data. P.L. analyzed the ribosome profiling data. L.S. and T.S. performed the transmission electron microscopy. T.S. and H.W. performed the mitochondria isolation and *in vitro* import assays. T.S. and C.M.H. wrote the manuscript.

## Author Information

Reprints and permissions information is available at www.nature.com/reprints. The authors declare no competing financial interests. Readers are welcome to comment on the online version of the paper. Correspondence and requests for materials should be addressed to C.M.H.

## Data deposition

The data reported in this paper have been deposited in the Gene Expression Omnibus (GEO) database, https://www.ncbi.nlm.nih.gov/geo (accession no. GSE114951).

## Notes

### Competing Interest Statement

The authors have declared no competing interest.

## References

1 Nargund, A. M., Pellegrino, M. W., Fiorese, C. J., Baker, B. M. & Haynes, C. M. Mitochondrial import efficiency of ATFS-1 regulates mitochondrial UPR activation. Science 337, 587–590, doi:10.1126/science.1223560 (2012).

2 Matthews, G. D., Gur, N., Koopman, W. J., Pines, O. & Vardimon, L. Weak mitochondrial targeting sequence determines tissue-specific subcellular localization of glutamine synthetase in liver and brain cells. J Cell Sci 123, 351–359, doi:10.1242/jcs.060749 (2010).

3 Saxton, R. A. & Sabatini, D. M. mTOR Signaling in Growth, Metabolism, and Disease. Cell 168, 960–976, doi:10.1016/j.cell.2017.02.004 (2017).

4 Baker, B. M., Nargund, A. M., Sun, T. & Haynes, C. M. Protective coupling of mitochondrial function and protein synthesis via the eIF2alpha kinase GCN-2. PLoS Genet 8, e1002760, doi:10.1371/journal.pgen.1002760 (2012).

5 Haynes, C. M., Petrova, K., Benedetti, C., Yang, Y. & Ron, D. ClpP mediates activation of a mitochondrial unfolded protein response in C. elegans. Dev Cell 13, 467–480, doi:10.1016/j.devcel.2007.07.016 (2007).

6 Rauthan, M., Ranji, P., Aguilera Pradenas, N., Pitot, C. & Pilon, M. The mitochondrial unfolded protein response activator ATFS-1 protects cells from inhibition of the mevalonate pathway. Proc Natl Acad Sci U S A 110, 5981–5986, doi:10.1073/pnas.1218778110 (2013).

7 Berendzen, K. M. et al. Neuroendocrine Coordination of Mitochondrial Stress Signaling and Proteostasis. Cell 166, 1553–1563 e1510, doi:10.1016/j.cell.2016.08.042 (2016).

8 Sorrentino, V. et al. Enhancing mitochondrial proteostasis reduces amyloid-beta proteotoxicity. Nature 552, 187–193, doi:10.1038/nature25143 (2017).

9 Haynes, C. M., Yang, Y., Blais, S. P., Neubert, T. A. & Ron, D. The matrix peptide exporter HAF-1 signals a mitochondrial UPR by activating the transcription factor ZC376.7 in C. elegans. Mol Cell 37, 529–540, doi:10.1016/j.molcel.2010.01.015 (2010).

10 Nargund, A. M., Fiorese, C. J., Pellegrino, M. W., Deng, P. & Haynes, C. M. Mitochondrial and nuclear accumulation of the transcription factor ATFS-1 promotes OXPHOS recovery during the UPR(mt). Mol Cell 58, 123–133, doi:10.1016/j.molcel.2015.02.008 (2015).

11 Pfanner, N., Warscheid, B. & Wiedemann, N. Mitochondrial proteins: from biogenesis to functional networks. Nat Rev Mol Cell Biol, doi:10.1038/s41580-018-0092-0 (2019).

12 Lin, Y. F. et al. Maintenance and propagation of a deleterious mitochondrial genome by the mitochondrial unfolded protein response. Nature 533, 416–419, doi:10.1038/nature17989 (2016).

13 Deng, P. et al. Mitochondrial UPR repression during Pseudomonas aeruginosa infection requires the bZIP protein ZIP-3. Proc Natl Acad Sci U S A, doi:10.1073/pnas.1817259116 (2019).

14 Narendra, D., Tanaka, A., Suen, D. F. & Youle, R. J. Parkin is recruited selectively to impaired mitochondria and promotes their autophagy. J Cell Biol 183, 795–803, doi:10.1083/jcb.200809125 (2008).

15 Fukasawa, Y. et al. MitoFates: improved prediction of mitochondrial targeting sequences and their cleavage sites. Mol Cell Proteomics 14, 1113–1126, doi:10.1074/mcp.M114.043083 (2015).

16 Melber, A. & Haynes, C. M. UPR(mt) regulation and output: a stress response mediated by mitochondrial-nuclear communication. Cell Res 28, 281–295, doi:10.1038/cr.2018.16 (2018).

17 Dillin, A., Crawford, D. K. & Kenyon, C. Timing requirements for insulin/IGF-1 signaling in C. elegans. Science 298, 830–834, doi:10.1126/science.1074240 (2002).

18 Zhang, Y. et al. Neuronal TORC1 modulates longevity via AMPK and cell nonautonomous regulation of mitochondrial dynamics in C. elegans. Elife 8, doi:10.7554/eLife.49158 (2019).

19 Gatsi, R. et al. Prohibitin-mediated lifespan and mitochondrial stress implicate SGK-1, insulin/IGF and mTORC2 in C. elegans. PLoS One 9, e107671, doi:10.1371/journal.pone.0107671 (2014).

20 Mair, W. et al. Lifespan extension induced by AMPK and calcineurin is mediated by CRTC-1 and CREB. Nature 470, 404–408, doi:10.1038/nature09706 (2011).

21 Schalm, S. S. & Blenis, J. Identification of a conserved motif required for mTOR signaling. Curr Biol 12, 632–639 (2002).

22 Land, S. C. & Tee, A. R. Hypoxia-inducible factor 1alpha is regulated by the mammalian target of rapamycin (mTOR) via an mTOR signaling motif. J Biol Chem 282, 20534–20543, doi:10.1074/jbc.M611782200 (2007).

23 Feng, J., Bussiere, F. & Hekimi, S. Mitochondrial electron transport is a key determinant of life span in Caenorhabditis elegans. Dev Cell 1, 633–644 (2001).

24 Labrousse, A. M., Zappaterra, M. D., Rube, D. A. & van der Bliek, A. M. C. elegans dynamin-related protein DRP-1 controls severing of the mitochondrial outer membrane. Mol Cell 4, 815–826, doi:10.1016/s1097-2765(00)80391-3 (1999).

25 Stadler, M., Artiles, K., Pak, J. & Fire, A. Contributions of mRNA abundance, ribosome loading, and post- or peri-translational effects to temporal repression of C. elegans heterochronic miRNA targets. Genome Res 22, 2418–2426, doi:10.1101/gr.136515.111 (2012).

26 Durieux, J., Wolff, S. & Dillin, A. The cell-non-autonomous nature of electron transport chain-mediated longevity. Cell 144, 79–91, doi:10.1016/j.cell.2010.12.016 (2011).

27 Ordureau, A. et al. Global Landscape and Dynamics of Parkin and USP30-Dependent Ubiquitylomes in iNeurons during Mitophagic Signaling. Mol Cell 77, 1124–1142 e1110, doi:10.1016/j.molcel.2019.11.013 (2020).

28 Phu, L. et al. Dynamic Regulation of Mitochondrial Import by the Ubiquitin System. Mol Cell 77, 1107–1123 e1110, doi:10.1016/j.molcel.2020.02.012 (2020).

29 Gass, J. N., Gunn, K. E., Sriburi, R. & Brewer, J. W. Stressed-out B cells? Plasma-cell differentiation and the unfolded protein response. Trends Immunol 25, 17–24, doi:10.1016/j.it.2003.11.004 (2004).

30 Yoneda, T. et al. Compartment-specific perturbation of protein handling activates genes encoding mitochondrial chaperones. J Cell Sci 117, 4055–4066, doi:10.1242/jcs.01275 (2004).

31 Benedetti, C., Haynes, C. M., Yang, Y., Harding, H. P. & Ron, D. Ubiquitin-like protein 5 positively regulates chaperone gene expression in the mitochondrial unfolded protein response. Genetics 174, 229–239, doi:10.1534/genetics.106.061580 (2006).

32 Paix, A., Folkmann, A., Rasoloson, D. & Seydoux, G. High Efficiency, Homology-Directed Genome Editing in Caenorhabditis elegans Using CRISPR-Cas9 Ribonucleoprotein Complexes. Genetics 201, 47–54, doi:10.1534/genetics.115.179382 (2015).

33 Dokshin, G. A., Ghanta, K. S., Piscopo, K. M. & Mello, C. C. Robust Genome Editing with Short Single-Stranded and Long, Partially Single-Stranded DNA Donors in Caenorhabditis elegans. Genetics 210, 781–787, doi:10.1534/genetics.118.301532 (2018).

34 Mello, C. C., Kramer, J. M., Stinchcomb, D. & Ambros, V. Efficient gene transfer in C.elegans: extrachromosomal maintenance and integration of transforming sequences. EMBO J 10, 3959–3970 (1991).

35 Rual, J. F. et al. Toward improving Caenorhabditis elegans phenome mapping with an ORFeome-based RNAi library. Genome Res 14, 2162–2168, doi:10.1101/gr.2505604 (2004).

36 Valenci, I., Yonai, L., Bar-Yaacov, D., Mishmar, D. & Ben-Zvi, A. Parkin modulates heteroplasmy of truncated mtDNA in Caenorhabditis elegans. Mitochondrion 20, 64–70, doi:10.1016/j.mito.2014.11.001 (2015).

37 Koopman, M. et al. A screening-based platform for the assessment of cellular respiration in Caenorhabditis elegans. Nat Protoc 11, 1798–1816, doi:10.1038/nprot.2016.106 (2016).

38 Robinson, M. D., McCarthy, D. J. & Smyth, G. K. edgeR: a Bioconductor package for differential expression analysis of digital gene expression data. Bioinformatics 26, 139–140, doi:10.1093/bioinformatics/btp616 (2010).

39 Schindelin, J. et al. Fiji: an open-source platform for biological-image analysis. Nat Methods 9, 676–682, doi:10.1038/nmeth.2019 (2012).

40 Grove, C. et al. Using WormBase: A Genome Biology Resource for Caenorhabditis elegans and Related Nematodes. Methods Mol Biol 1757, 399–470, doi:10.1007/978-1-4939-7737-6_14 (2018).

41 Subramanian, A. et al. Gene set enrichment analysis: a knowledge-based approach for interpreting genome-wide expression profiles. Proc Natl Acad Sci U S A 102, 15545–15550, doi:10.1073/pnas.0506580102 (2005).

42 Wingett, S. W. & Andrews, S. FastQ Screen: A tool for multi-genome mapping and quality control. F1000Res 7, 1338, doi:10.12688/f1000research.15931.2 (2018).

43 Blankenberg, D. et al. Manipulation of FASTQ data with Galaxy. Bioinformatics 26, 1783–1785, doi:10.1093/bioinformatics/btq281 (2010).

44 Ihaka, R. & Gentleman, R. R: A Language for Data Analysis and Graphics. Journal of Computational and Graphical Statistics 5, 299–314 (1996).

45 Weidberg, H. & Amon, A. MitoCPR-A surveillance pathway that protects mitochondria in response to protein import stress. Science 360, doi:10.1126/science.aan4146 (2018).

