## Extended data for "UPR^mt^ scales mitochondrial network expansion with protein synthesis via mitochondrial import"

**Extended Data Figure 1**

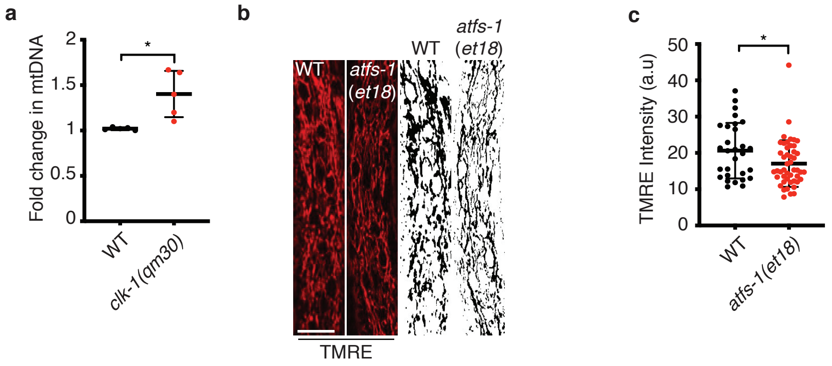

**Extended Data Figure 1. *clk-1(qm30)* harbor more mtDNAs than wildtype worms.**

**a.** Quantification of mtDNA in wildtype, and *clk-1(qm30)* worms as determined by qPCR. N=4. Error bars mean +/- s.d, *p<0.05 (Student's *t*-test).

**b.** TMRE staining of wildtype and *atfs-1(et18)* worms. Scale bar 10 μm.

**c.** Quantification of TMRE intensity of panel 1b. n=29 (wildtype) n=47 (*atfs-1(et18)*), Error bars mean +/- s.d, *p<0.05.

**Extended Data Figure 2**

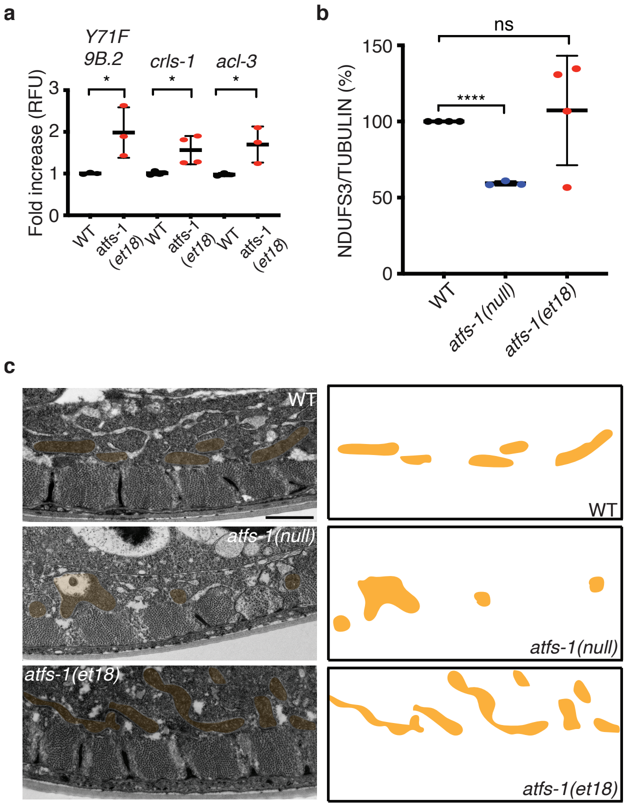

**Extended Data Figure 2. mRNAs encoding for mitochondrial proteins are differentially expressed in *atfs-1(et18)* and *atfs-1(null)* worms relative to wildtype worms.**

**a.** Transcript levels of the TAM41 mitochondrial translocator assembly and maintenance homolog (*Y71F9B.2*), acyltransferase-like 3 (*acl-3*) and cardiolipin synthase homolog (*crls-1*) as determined by qRT-PCR in wildtype and *atfs-1(et18)* worms. N=3 or N=4 for *crls-1*. Error bars mean +/- s.d, *p<0.05 (Student's t-test).

**b.** Quantification of NDUFS3 levels relative to TUBULIN in *pdr-1(tm598)*, atfs-*1(null);pdr-1(tm598)* and *atfs-1(et18);pdr-1(tm598)* strains. N=3 *(atfs-1(null))*, N=4 *(wildtype and atfs-1(et18))*. Error bars mean +/- s.d, ***p<0.001 (Student's t-test).

**c.** Transmission electron microscopy of body wall muscles of wildtype, *atfs-1(null)* and *atfs-1(et18)*. Mitochondria are highlighted in yellow. Scale bar 1 μm.

**Extended Data Figure 3**

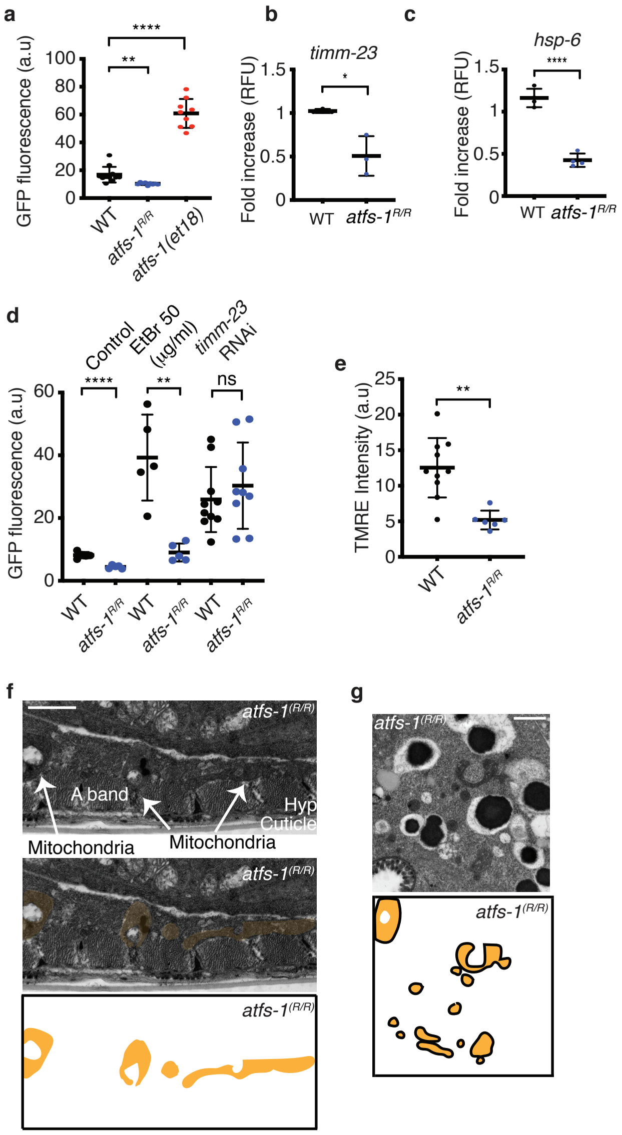

**Extended Data Figure 3. Increasing the strength of the ATFS-1 MTS inhibits the UPR$^{mt}$ and perturbs the mitochondrial network.**

**a.** Quantification of GFP intensity in wildtype, *atfs-1$^{R/R}$* and *atfs-1(et18)* worms expressing *hsp-6$_{pr}$::gfp*. N=11(wildtype), N=9(*atfs-1$^{R/R}$* and *atfs-1(et18)*). Error bars mean +/- s.d, **p<0.01,****p<0.0001 (Student's t-test).

**b-c.** Transcript levels of translocase of inner mitochondrial membrane-23 (*timm-23*) (**b**) and of heat shock protein-6 (*hsp-6*) (**c**) as determined by qRT-PCR in wildtype and *atfs-1$^{R/R}$* worms. N=4 (*hsp-6*), N=3 (*timm-23*). Error bars mean +/- s.d, *p<0.05, ****p<0.0001 (Student's t-test).

**d.** Quantification of GFP intensity in wildtype and *atfs-1$^{R/R}$* worms expressing *hsp-6$_{pr}$::gfp* and raised on control(RNAi), *timm-23*(RNAi) or 50 µg/ml EtBr. N=5 control and EtBr. For *timm-23*(RNAi) N=10,9(wildtype and *atfs-1$^{R/R}$*, respectively). Error bars mean +/- s.d, **p<0.01,****p<0.0001 (Student's t-test).

**e.** Quantification of TMRE intensity in wildtype and *atfs-1$^{R/R}$* worms. N=10,6 (wildtype and *atfs-1$^{R/R}$*, respectively). Error bars mean +/- s.d, **p<0.01 (Student's *t*-test).

**f.** Transmission electron microscopy of body wall muscle cells of *atfs-1$^{R/R}$* worms (wildtype control in Figure 2i). Mitochondria are highlighted in yellow**.** Scale bar 1 µm.

**g.** Transmission electron microscopy of intestinal cells of *atfs-1$^{R/R}$* worms. Mitochondria are highlighted in yellow. (wildtype control in Figure 2l). Scale bar 1 µm.

### Extended Data Figure 4

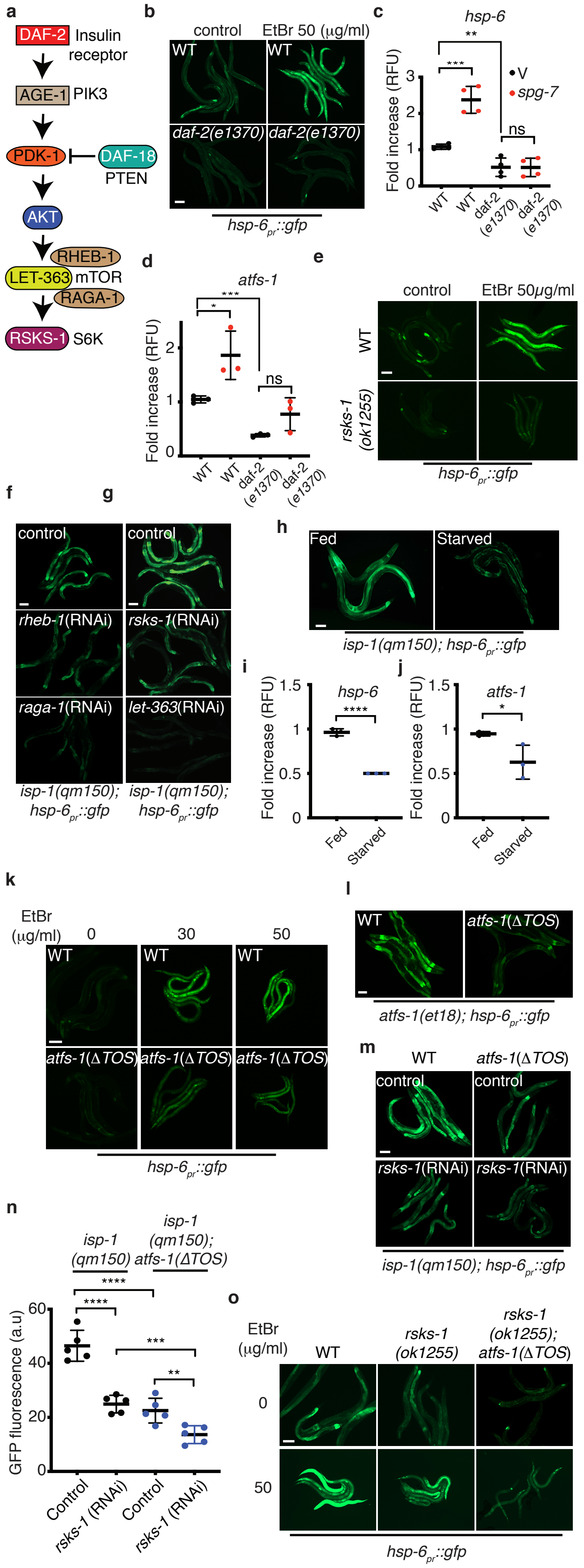

**Extended Data Figure 4. TORC1 regulates the UPR$^{mt}$**

**a.** Schematic of the *C. elegans* insulin-like signaling and TORC1 pathway. Mammalian homologs included. The small GTPase Rheb is required for TORC1 activity, while the GTPase RAGA-1 regulates TORC1 activity in response to amino acid availability.

**b.** *hsp-6$_{pr}$::gfp* and *daf-2(e1370);hsp-6$_{pr}$::gfp* worms raised on control or 50 µg/ml EtBr at 20°C. Scale bar 0.1 mm.

**c-d.** Transcript levels of heat shock protein-6 (*hsp-6*) (**c**) and activated transcription factor stress-1 (*atfs-1*) (**d**) as determined by qRT-PCR in wildtype and *daf-2(e1370)* strains raised on control(RNAi) or *spg-7*(RNAi) at 20°C. N=4(*hsp-6*), N=3(*atfs-1*) Error bars mean +/- s.d. *p<0.05, **p<0.01, ***p<0.001 (Student's t-test).

**e.** Photomicrographs of *hsp-6$_{pr}$::gfp* and *rsks-1(ok1255);hsp-6$_{pr}$::gfp* worms raised on 0 or 50 µg/ml EtBr. Scale bar 0.1 mm.

**f.** Photomicrographs of *isp-1(qm150)*;*hsp-6$_{pr}$::gfp* worms raised on control, *rheb-1* or *raga-1*(RNAi). Scale bar 0.1 mm.

**g.** Photomicrographs of *isp-1(qm150)*;*hsp-6$_{pr}$::gfp* worms raised on control, *rsks-1* or *let-363*(RNAi). Scale bar 0.1 mm.

**h.** Photomicrographs of *isp-1(qm150)*;*hsp-6$_{pr}$::gfp* worms raised to the L4 stage and starved for 1h. Scale bar 0.1 mm.

**i-j.** Transcript levels of heat shock protein-6 (*hsp-6*) (**i**) and activated transcription factor stress-1 (*atfs-1*) (**j**) as determined by qRT-PCR in worms raised to L4 and starved for 1h. Error bars mean +/- s.d. *p<0.05, ****p<0.0001 (Student's t-test).

**k.** *hsp-6$_{pr}$::gfp* and *atfs-1(∆TOS);hsp-6$_{pr}$::gfp* worms raised on 0,30 or 50 µg/ml EtBr. Scale bar 0.1 mm.

**l.** *atfs-1(et18)*;*hsp-6*$_{pr}$*::gfp* and *atfs-1(et18,∆TOS)*;*hsp-6*$_{pr}$*::gfp* worms. Scale bar 0.1mm.

**m.** *isp-1(qm150)*;*hsp-6*$_{pr}$*::gfp* and *isp-1(qm150)*;*atfs-1(∆TOS)*;*hsp-6*$_{pr}$*::gfp* worms raised on control or *rsks-1*(RNAi). Scale bar 0.1 mm.

**n.** Quantification of the experiment in panel m. N=5. Error bars mean +/- s.d, **p<0.01, ***p<0.001 ****p<0.0001 (Student's t-test).

**o.** *hsp-6*$_{pr}$*::gfp, rsks-1(ok1255);hsp-6*$_{pr}$*::gfp* and *rsks-1(ok1255);atfs-1(∆TOS);hsp-6*$_{pr}$*::gfp* worms raised on 0 or 50 µg/ml EtBr. Scale bar 0.1 mm.

**Extended Data Figure 5**

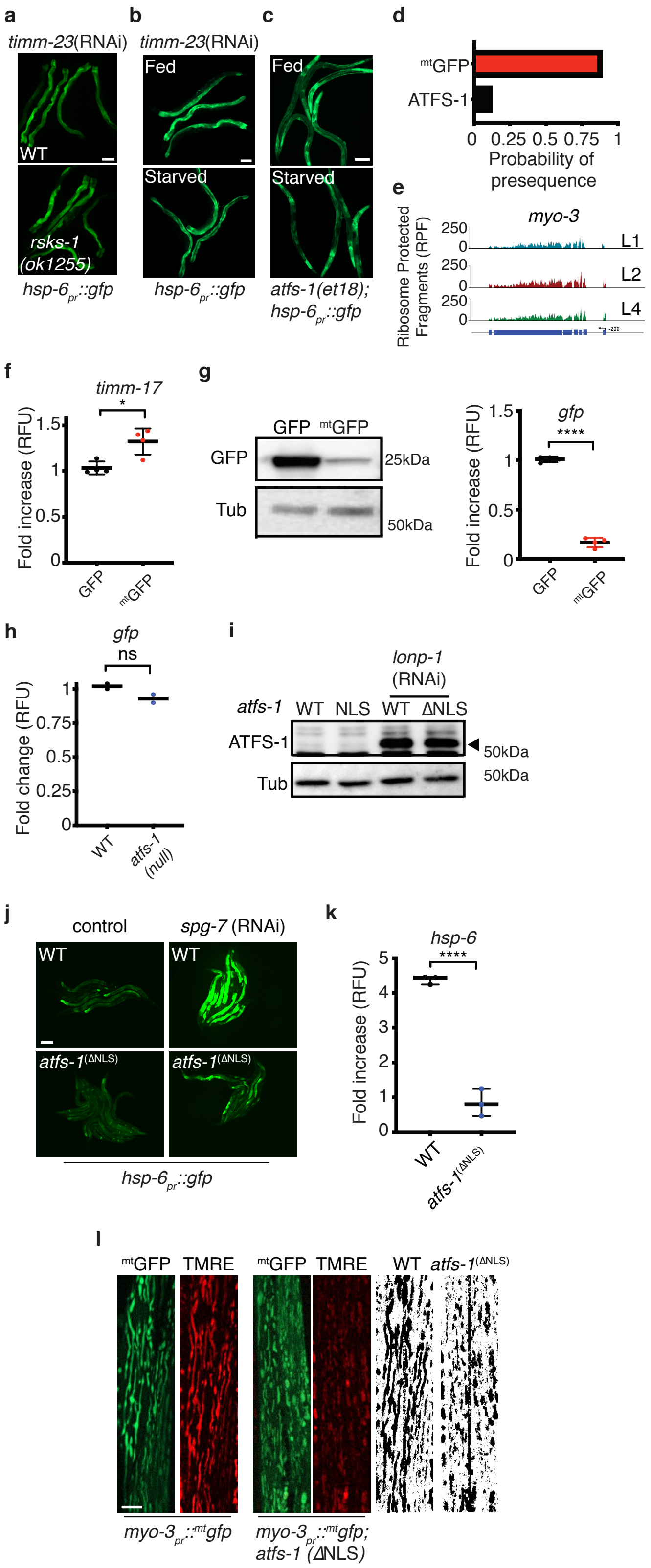

**Extended Data Figure 5. Mitochondrial expansion is regulated by mitochondrial protein import**

**a.** Photomicrographs of wildtype and *rsks-1(ok1255);hsp-6$_{pr}$::gfp* worms raised on *timm-23*(RNAi). Scale bar 0.1 mm.

**b.** Photomicrographs of *hsp-6$_{pr}$::gfp* worms raised on *timm-23*(RNAi) to the L4 stage and starved for 24 h. Scale bar 0.1 mm.

**c.** Photomicrographs of *atfs-1(et18)*;*hsp-6$_{pr}$::gfp* worms raised to the L4 stage and starved for 24 h. Scale bar 0.1 mm.

**d.** Mitochondrial targeting sequence probability prediction using MitoFates. The MTS of aspartate aminotransferase (amino acids 1-24) (AST) fused to GFP (red) and ATFS-1 (black) are presented.

**e.** Ribosome protected fragments (RPF) profile of *myo-3* at the L1, L2, and L4 larval developmental stages.

**f.** Transcript levels of translocase of inner mitochondrial membrane-17 (*timm-17*) as determined by qRT-PCR in *myo-3$_{pr}$::gfp* and in *myo-3$_{pr}$::$^{mt}$gfp* worms. N=4. Error bars mean +/- s.d, *p<0.05 (Student's t-test).

**g.** Immunoblots of GFP and Tubulin in *myo-3$_{pr}$::gfp* and *myo-3$_{pr}$::$^{mt}$gfp* expressing worms (left side) and transcript levels of green fluorescent protein (*gfp*) as determined by qRT-PCR in *myo-3$_{pr}$::gfp* and *myo-3$_{pr}$::$^{mt}$gfp* worms. N=4 Error bars mean +/- s.d (Student's t-test) ****p<0.0001 (right side).

**h.** Transcript levels of green fluorescent protein (*gfp*) as determined by qRT-PCR in *myo-3$_{pr}$::$^{mt}$gfp* and in *atfs-1(null)*;*myo-3$_{pr}$::$^{mt}$gfp* worms. N=2 Error bars mean +/- s.d (Student's t-test).

**i.** Immunoblots of wildtype and *atfs-1*$^{(\Delta NLS)}$ worms raised on control or *lonp-1*(RNAi). ATFS-1 (▸).

**j.** *hsp-6*$_{pr}$*::gfp* and *atfs-1*$^{(\Delta NLS)}$*;hsp-6*$_{pr}$*::gfp* worms raised on control or *spg-7*(RNAi). Scale bar 0.1 mm.

**k.** Transcript levels of heat shock protein-6 (*hsp-6*) as determined by qRT-PCR in wildtype and *atfs-1*$^{(\Delta NLS)}$ worms raised on *spg-7*(RNAi). N=3. Error bars mean +/- s.d. ****p<0.0001 (Student's t-test).

**l.** TMRE staining of wildtype and *atfs-1*$^{(\Delta NLS)}$ worms expressing *myo-3*$_{pr}$*::*$^{mt}$*gfp*. Scale bar 10 µm. Skeleton-like binary backbone are presented to the right.

**Extended Data Figure 6**

**a**

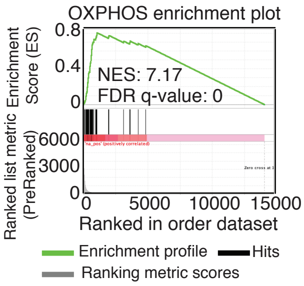


**b**

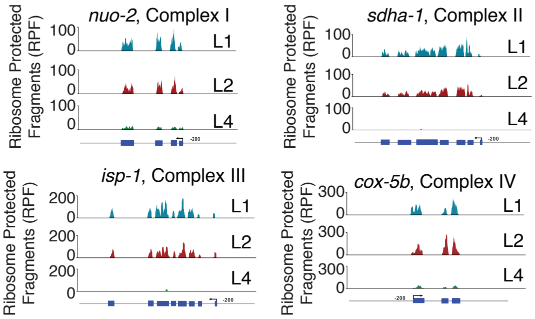


**c**

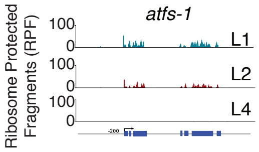

**Extended Data Figure 6. Translation of OXPHOS complex components primarily occurs during the early stages of worm development, similar to ATFS-1.**

**a.** Gene Set Enrichment Analysis (GSEA) comparing the expression of OXPHOS genes to all other genes in *C. elegans*. Gene expression abundances were measured and ranked by reads per kilobase per million reads (RPKM) from the RNA-seq data of WT L4 worms. Each black line represents an OXPHOS gene and each white line represents genes that are not OXPHOS genes.

**b.** Ribosome protected fragments (RPF) profile of four representative OXPHOS complex subunits (*nuo-2* – complex I, *sdha-1* – complex 2, *isp-1* – complex 3 and *cox-5b* -complex 4) at different larval developmental stages (L1, L2 and L4).

**c.** Ribosome protected fragments (RPF) profile of *atfs-1* at the L1, L2, and L4 larval developmental stages.
